# Hypertriglyceridemia and obesity exacerbate the course of SIRS induced by SAP in Rats

**DOI:** 10.1101/663104

**Authors:** Kelei Hua, RuiXia Li, LiYing Cao, WanSheng Lao

**Affiliations:** Department of Surgery, The Affiliated Cancer Hospital of Zhengzhou University, Henan Provincial Cancer Hospital, Zhengzhou, Henan450000, P.R. China; Department of Surgery, the People’s Hospital of Jiaozuo City, Jiaozuo, Henan 454000, P.R. China; Department of Surgery, The Affiliated Tangshan Kailuan hospital of North China University of Science and Technology, Tangshan, Hebei063000, P.R. China; Department of Surgery, Yangxin county hospital of traditional Chinese medicine, Binzhou256600, Shandong, P.R. China

**Keywords:** Hypertriglyceridaemia, Obesity, Systemic Inflammatory Response Syndrome, Pancreatitis, Acute Necrotizing

## Abstract

The aim of the present study was to explore the mechanism underlying how HTG (hypertriglyceridaemia) and obesity exacerbate the course of the systemic inflammatory response syndrome (SIRS) induced by severe acute pancreatitis (SAP) in rats. Seventy-two rats were fed a normal or high-fat diet to induce HTG and obesity, and SAP was induced by retrograde injection of 5% sodium taurocholate solution at a volume of 1 ml/kg into the biliopancreatic duct. The injury to the pancreas was assessed by macroscopic observation, pancreatic histological evaluation and serum levels of amylase and lipase. SIRS was estimated by measuring SIRS scores and interleukin-6 (IL-6), tumour necrosis factor alpha (TNF-α) and interleukin-10 (IL-10) expression. The results showed that the SIRS scores and pancreatic histological scores increased significantly and the blood calcium level decreased significantly in the hypertriglyceridaemia SAP (HSAP) group compared with those of the SAP group. In addition, HTG and obesity significantly increased plasma levels of the proinflammatory cytokines IL-6 and TNF-α and significantly downregulated the proinflammatory cytokine IL-10. Our findings showed that HSAP rats exhibited more severe pancreatic injury and more serious SIRS scores than the SAP rats did. The underlying mechanism may be that HTG and obesity intensify early-stage SIRS by regulating the levels of inflammatory and anti-inflammatory cytokines.

## Introduction

Acute pancreatitis (AP) is a frequent inflammatory disease of the pancreas with multiple causes, among which hypertriglyceridemia is the third most common and accounts for up to approximately 10% of all cases (1). Clinical studies have reported that SAP patients with HTG and obesity suffer a more severe clinical course and complications, including aggravating systemic inflammatory response,multiple organ failure, infection and high mortality(2–4). Animal experiments have also provided evidence that HTG with SAP could exacerbate oedema and necrosis of the pancreas (5). However, the exact mechanism remains unclear.

SIRS is an early manifestation of multiple organ dysfunction syndrome (MODS) and multiple organ failure (MOF)(6). SAP has two peak time points at which the risk of mortality is high: the first is early MODS, which is usually caused by the continued deterioration of SIRS, and the second occurs as a result of sepsis(7). Therefore, in the early stage of pancreatitis, any factor aggravating SIRS will increase the mortality rate of patients. Based on the above knowledge, we hypothesized that HTG and obesity may play an important role in the duration and severity of early-stage SIRS. Therefore, this study was performed to explore the mechanism underlying how HTG and obesity intensify the course of SIRS induced by SAP in rats.

## Materials and Methods

### Experimental animals and protocols

All animal experimental protocols were approved by the Animal Care and Use Committee of the North China University of Science and Technology, and the experiments were performed in accordance with the guidelines of the committee (approval number: 20130051).

Male Sprague-Dawley rats weighing 110–130 g were purchased from Beijing HFK Bioscience Co. Ltd. (Beijing, China). All rats were housed under controlled day-night cycles. The experimental design is shown in Figure 1. A total of 72 SD rats were randomly allocated to two groups (normal lipids and HTG, n=36 in each group). The normal lipid group was divided into sham-operated (SO) and SAP groups with 18 rats in each group. Each group was then divided into 2 h, 6 h and 24 h subgroups. The HTG group was divided into SO and SAP groups with 18 rats in each group. Each group was then divided into 2 h, 6 h and 24 h subgroups. HTG rats were fed a high-fat diet (HFD) (D12492 diet formulation, protein 20%, carbohydrate 20%, fat 60%), and the normal lipid rats were fed a normal diet. After 8 weeks of HFD feeding, SAP groups were induced by retrograde injection of 5% sodium taurocholate solution at a volume of 1 ml/kg into the biliopancreatic duct. The SO groups had only their abdomens opened and closed. At each of the desired time points (2 h, 6 h and 24 h after SAP induction), intraperitoneal injections of 10% hydrate chloral (0.3 ml/100 g) were used for surgical anaesthesia. Heart rate was measured with a tail cuff (BIOPAC MP150, USA), temperature was measured via the anus (BIOPAC MP150, USA), and blood samples were taken from the abdominal aorta. PaCO_2_ was measured in the samples using a blood gas analyser (Radiometer, ABL500, Kongeriget, Denmark), and white cells were counted using a haematology analyser (SYSMEX, SysmexXT-1800i, Japan). The pancreas was promptly fixed in 10% neutral buffered formaldehyde solution for further histological examination. SIRS criteria and score calculations were carried out following the description of Lu(8). Briefly, one point was recorded for a heart rate increase of 50% above the normal range, a 2-fold increase or 50% decrease in the peripheral total white blood cell count compared with normal ranges, a core temperature increase or decrease of 1°C relative to the normal range, and a PaCO_2_ decrease of 25% below the normal range. The scoring range was between 0 and 4. When the score was 2 or more, the rat was defined to be in a state of SIRS.

**Figure 1:**
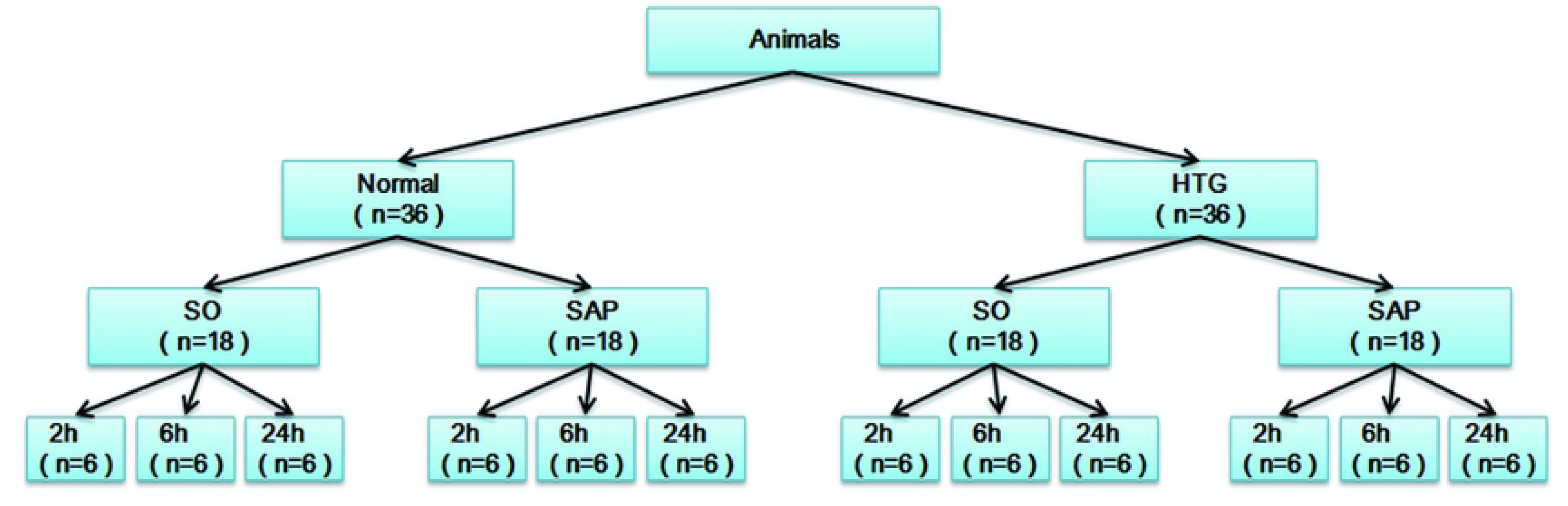
Study design. SO, sham-operated; SAP, severe acute pancreatitis; SO2, 6, 24 h, normal lipid + sham-operated + sacrificed at 2, 6, or 24 h; SAP2, 6, 24 h, normalm lipid + SAP + sacrificed at 2, 6, or 24 h; HSO2, 6, 24 h, HTG + sham-operated + sacrificed at 2, 6, or 24 h; HSAP2, 6, 24 h, HTG + SAP + sacrificed at 2, 6, or 24 h.

### Serum biochemistry

Serum levels of amylase, lipase, calcium (Ca2+), triglycerides (TGs) and total cholesterol (TC) were analysed using an automated biochemical analyser (7600, Hitachi, Japan).

### Analysis of TNF-a, IL-6, and IL-10 Levels in the Serum

Briefly, blood samples were collected and clotted at room temperature, and serum samples were obtained following centrifugation. Serum TNF-α, IL-6 and IL-10 levels were assessed using ELISA kits (Cat# 88-7324-22, BMS603-2, 88-7105-22 Thermofisher, USA). The intensity of the colour was measured using a microplate reader (Thermal, USA).

### Evaluation of Pancreatic Pathologies

The pancreas of each rat was fixed with 10% formaldehyde, embedded in paraffin, cut into 4 μm sections and stained with haematoxylin and eosin. The scoring system applied to evaluate the histological examinations followed a previous description(9, 10), as shown in Table 1. The evaluation was performed by two investigators who were blind to the experimental treatment. Five fields in each sample were observed, and the total scores of the five fields were averaged.

**Table 1:** Histological scoring for SAP

|  | Score | Description |
| --- | --- | --- |
| Edema | 0 | Absent |
|  | 1 | Diffuse expansion of interlobar septa |
|  | 2 | 1+ diffuse expansion of interlobular septa |
|  | 3 | 2+ diffuse expansion of interacinar septa |
|  | 4 | 3+ diffuse expansion of intercellular septa |
| Inflammation | 0 | Absent |
|  | 1 | Around ductal margins |
| | 2 | In parenchyma ($\leq 50\%$ of lobules) |
|  | 3 | In parenchyma (51–75% of lobules) |
| | 4 | In parenchyma ($>75\%$ of lobules) |
| Hemorrhage | 0 | Absent |
| | 1 | Blood in parenchyma ($\leq 25\%$) |
|  | 2 | Blood in parenchyma (26–50%) |
|  | 3 | Blood in parenchyma (51–75%) |
| | 4 | Blood in parenchyma ($>75\%$) |
| Necrosis | 0 | Absent |
|  | 1 | Periductal parenchymal destruction |
| | 2 | Focal parenchymal necrosis ($\leq 20\%$) |
|  | 3 | Diffuse loss of lobules (21–50%) |
| | 4 | Severe loss of lobules ($>50\%$) |

### Statistical analysis

SPSS 21.0 was used for statistical analyses. The SIRS scores are expressed as the mean ± 95% CI and were assessed using nonparametric tests. Other data are expressed as the mean ± standard deviation. Unpaired Student’s t-tests were used for two-group comparisons, and one-way ANOVAs were used for multigroup comparisons. A significant difference was accepted if *P* < 0.05.

## Results

### Clinical manifestations

Rats were fed a HFD for 8 weeks to establish the HTG and obesity model. The mean body weight and visceral fat of the HFD group increased significantly compared to those of the normal diet group (*p<*0.001; Fig. 2A, C). The serum TG level was more than 3-fold higher in the HFD group than that in the normal diet group (*p<*0.0001). The TC level was also significantly higher (*p*<0.0001; Fig. 2B); following the injection of 5% sodium taurocholate solution, rats exhibited rapid breathing, and symptoms (heart rate, temperature, PaCO_2_ and white cells) were aggravated for a prolonged time. Compared with the SO group, plasma amylase and lipase levels were much higher in the SAP group, with statistically significant differences (*p*=0.021; Fig. 5A, B), thus confirming that the HTG and obesity SAP model was established.

**Figure 2:**
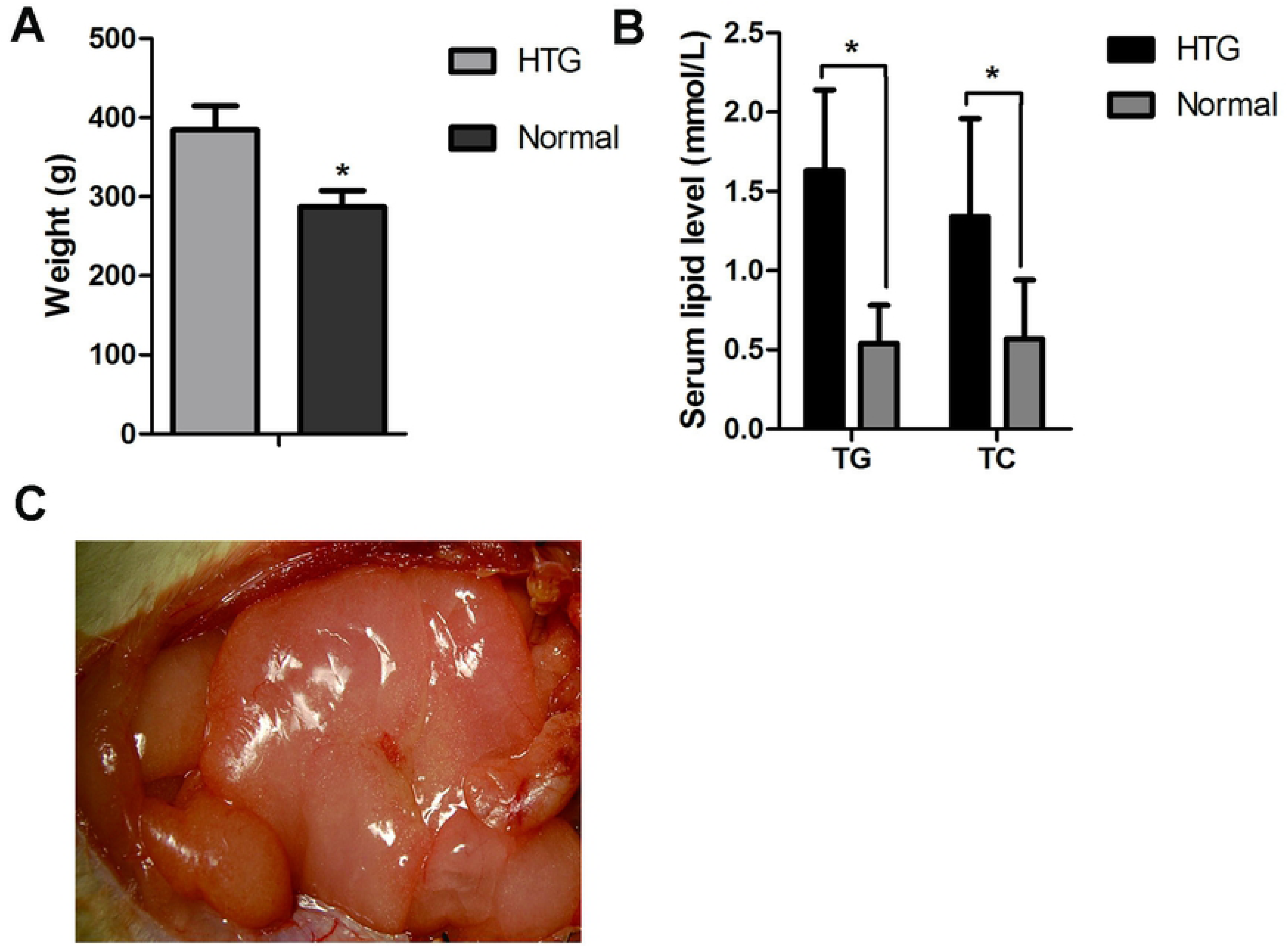
Effects of a high-fat diet on rats. A. Bar graph of body weight in normal lipid and HTG groups. B. Bar graph of serum TG and TC levels in normal lipid and HTG groups. C. Visceral fat in the HTG group.

### HTG and obesity exacerbate the course of SIRS

The SIRS scores from each group were calculated based on the above criteria. As shown in Figure 3C, the SIRS scores were notably higher in the SAP group than in the SO group. Compared with those in the SAP group, the SIRS scores increased in the HSAP group, especially at the 24 h time-point. We also found that the plasma levels of the proinflammatory cytokines IL-6 and TNF-a were markedly higher in the HSAP group than in the SAP group and that the down-regulation of the proinflammatory cytokine IL-10 was significantly decreased in the HSAP group compared to that in the SAP group, and all were significantly different at the 24 h time point (Fig. 5B, C, E).

**Figure 3:**
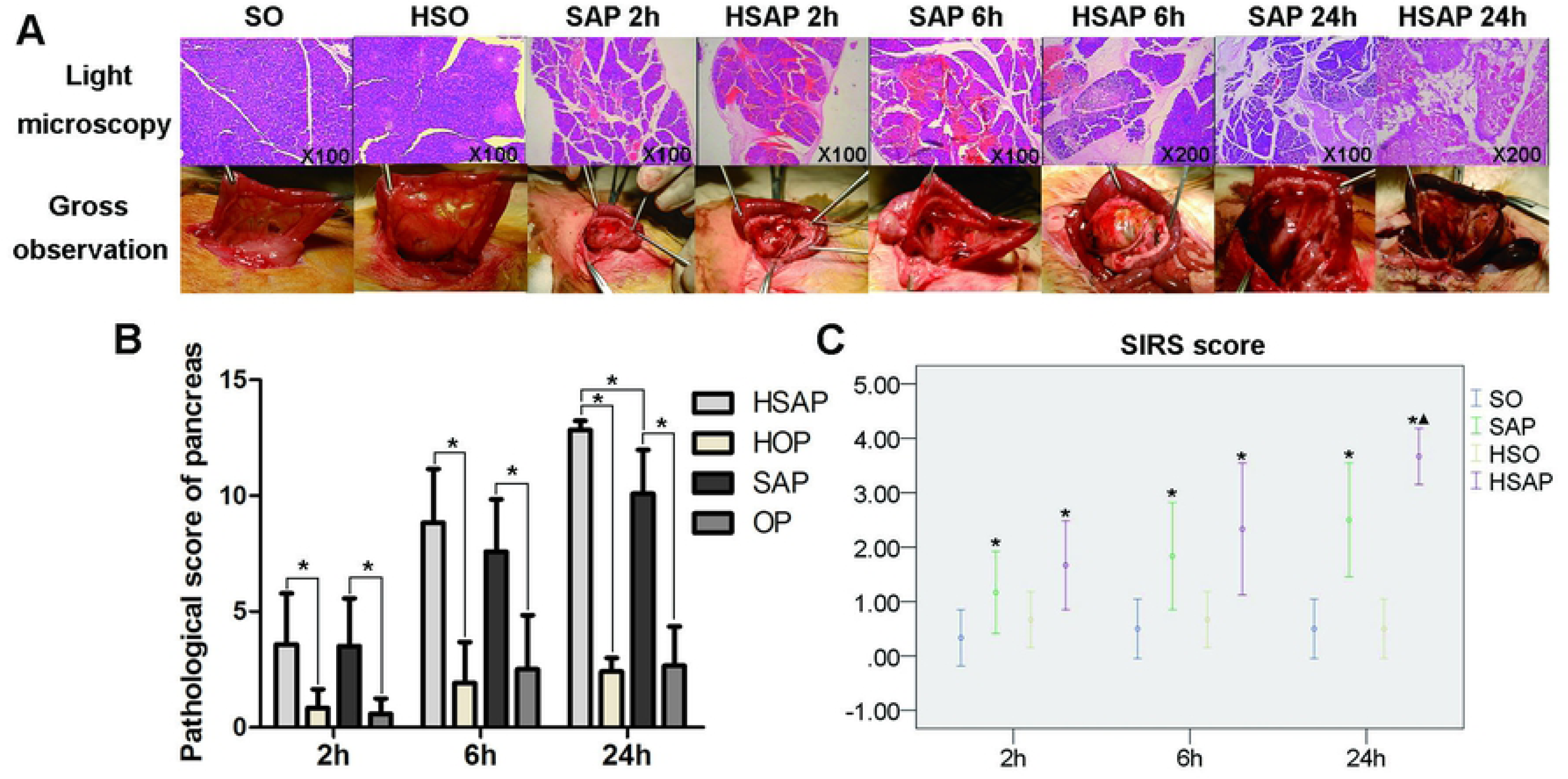
HTG and obesity aggravated pathological changes in the pancreas and SIRS scores. A. Gross and microscopic observation (HE×200) of pancreatic tissue at different times. B. The pancreatic pathological scores in the SAP group were notably high in the 24 h group* P< 0.05. C. The SIRS scores were notably high in the HSAP 24 h group. *Compared with SO groups P< 0.05, ▲ compared with SAP groups P< 0.05.

### HTG and obesity aggravated Pathological Changes in the Pancreas

In all SAP rats, the pancreatic injuries were featured by oedema, inflammation, haemorrhage, and patchy necrosis. The injuries in the pancreas had higher histopathological scores in the HSAP groups than in the SAP groups in the 24 h groups (*p* <0.001 Fig. 3B). We also observed gross observational changes in the pancreas, including evident regional or diffuse hyperaemia and oedema, with increased pancreatic envelope tension. After 6 h or 24 h, pancreatic haemorrhaging, necrosis and bloody ascites were observed in the surviving rats of the model groups. Consistent with the histopathology of the pancreas, the gross observational changes in the HSAP groups were also more severe compared with those of the NSAP groups (*p*=0.014). However, in the SO group, mild oedema of the pancreas and exudation in the abdominal cavity were observed (Fig. 3A).

### HTG and obesity aggravated hypocalcaemia

As shown in Figure 4B, the blood calcium level of the SAP group, compared with that of the SO group, significantly decreased after SAP induction (*p<*0.05). Compared with that in the SAP groups, blood calcium levels decreased significantly in the HSAP groups at the 24 h time point (*p*=0.003). Consistent with the above results, we also observed more saponification spots on the small bowel mesentery in the HSAP groups (Fig. 4A).

**Figure 4:**
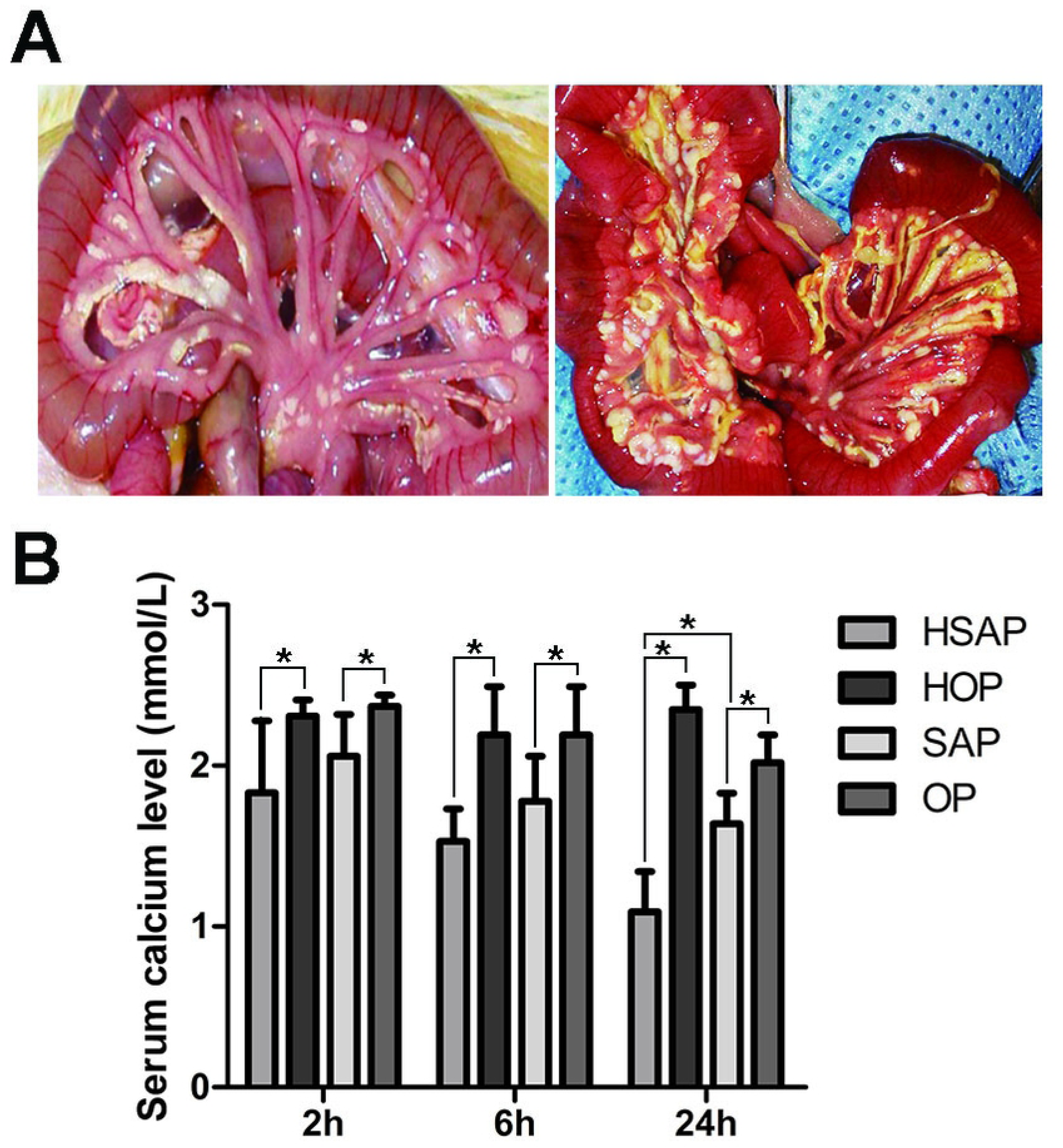
HTG and obesity aggravate hypocalcaemia. A. Saponification on the mesentery of the small intestine B. Bar graph of serum calcium.

**Figure 5:**
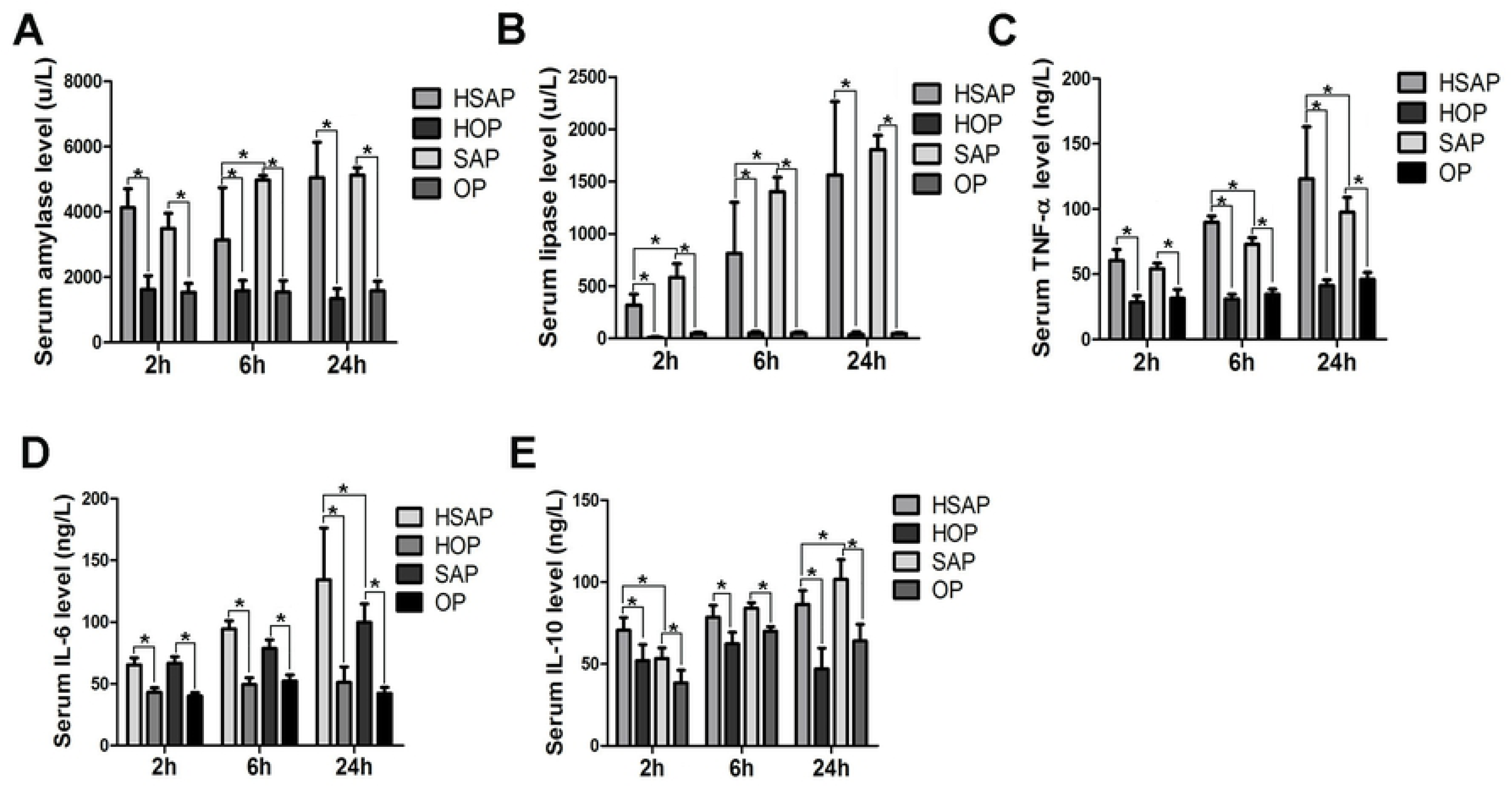
Effects of HTG and obesity on serum amylase, lipase, TNF-α, IL-6 and IL- 10. A, B, C, D, E: Bar graphs of serum amylase, lipase, TNF-α, IL-6 and IL-10, respectively.

## Discussion

Compared with the main aetiological factors, such as gallstones and alcohol abuse, HTG is the third most important factor in SIRS, being found in up to 7% of cases(11). In addition, HTG pancreatitis tends to be more severe than other causes(12). Previous clinical research has shown that persistent SIRS in HTG patients was higher than that in normal patients(13). Moreover, obesity likely contributes to increasing the incidence and severity of acute pancreatitis(14) and could serve as a prognostic indicator(15). However, the exact mechanism remains unclear.

In our study, we note that the pathological scores in the HSAP group were significantly higher than those in the SAP group at 24 h, which was similar to the results obtained in previous studies(5, 16). The SIRS score of the HSAP group was also significantly higher than that of the SAP group, especially at the 24 h time-point. These results indicate that HTG and obesity can continue to exacerbate SIRS over time. It is well known that multiple organ failure in patients with SAP is mainly due to aggravated SIRS, so any factor that aggravates SIRS affects the prognosis of patients with SAP. TNF-α is produced by activated macrophages. Its concentration correlates positively with the severity of pancreatic damage and SIRS(17). IL-6 is derived from a wide range of cells; the level of IL-6 rises in patients with AP, and its level correlates with the severity of AP (18). IL-10 is an anti-inflammatory pleiotropic cytokine that inhibits the inflammatory immune response through various mechanisms and has been demonstrated to inhibit the expression of TNF-α, IL-1, IL-6, IL-8, and IL-12 by monocytes and macrophages. We also note that the proinflammatory cytokines TNF-a and IL-6 were increased and the anti-inflammatory cytokine IL-10 was decreased in the HSAP group compared with the SAP group. These results confirm that HTG and obesity intensify early-stage SIRS by regulating the levels of inflammatory and anti-inflammatory cytokines. Unlike subcutaneous fat, which rarely impacts the severity of acute pancreatitis(19), obesity-associated increases in visceral fat can worsen acute pancreatitis outcomes(20) and convert mild AP to SAP in animal studies(21). In our study, visceral fat increased after 8 weeks of a high-fat diet.

Hypocalcaemia is one of the components of Ranson’s scoring system that is used to assess the severity of pancreatitis. Previous studies in animal models have shown that hypocalcaemia is a marker of poor prognosis in patients with pancreatitis(22). The mechanisms proposed for hypocalcaemia are autodigestion of mesenteric fat by pancreatic enzymes and release of free fatty acids, which undergo saponification. In our study, blood calcium levels decreased significantly in the HSAP groups at the 24 h time-point, and the macroscopic observation of adipose tissue showed that the saponification on the mesentery of the small intestine in the HSAP group was greater than that in the SAP group.

## Conclusion

Our findings showed that HSAP rats had more severe pancreatic injury and more serious SIRS scores than the SAP rats did. The underlying mechanism may be the intensification of early-stage SIRS related to HTG and obesity as a result of the imbalance of inflammatory and anti-inflammatory cytokines.

## Acknowledgements

The authors would like to thank Dr. Rui Li for his valuable suggestions.

## Conflicts of interest

The authors declare that they have no conflicts of interest

## Ethical approval

All procedures performed in the studies were in accordance with the ethical standards of the institution at which the studies were conducted.

